# Working memory predicts long-term recognition of auditory sequences: Dissociation between confirmed predictions and prediction errors

**DOI:** 10.1101/2024.09.20.614110

**Authors:** L. Bonetti, E. Risgaard Olsen, F. Carlomagno, E. Serra, S.A. Szabó, M. Klarlund, M.H. Andersen, L. Frausing, P. Vuust, E. Brattico, M.L. Kringelbach, G. Fernández-Rubio

## Abstract

Memory is a crucial cognitive process involving several subsystems: sensory memory (SM), short-term memory (STM), working memory (WM), and long-term memory (LTM). While each has been extensively studied, the interaction between WM and LTM, particularly in relation to predicting temporal sequences, remains largely unexplored. This study investigates the relationship between WM and LTM, and how these relate to aging and musical training. Using three datasets with a total of 244 healthy volunteers across various age groups, we examined the impact of WM on LTM recognition of novel and previously memorized musical sequences. Our results show that WM abilities are significantly related to recognition of novel sequences, with a more pronounced effect in older compared to younger adults. In contrast, WM did not similarly impact the recognition of memorized sequences, which implies that different cognitive processes are involved in handling prediction errors compared to confirmatory predictions, and that WM contributes to these processes differently. Additionally, our findings confirm that musical training enhances memory performance. Future research should extend our investigation to populations with cognitive impairments and explore the underlying neural substrates.

## Introduction

Memory is a fundamental cognitive process that enables the storage, retrieval, and use of information—such as ideas, events, stimuli, skills, and images—once the original input is no longer present ^1–5^. Memory is typically divided into three key subsystems, each serving distinct functions while interacting with the environment. These subsystems are (i) sensory memory (SM) ^6,7^ and short-term memory (STM) ^8,9^, (ii) working memory (WM) ^10–12^, and (iii) long-term memory (LTM) ^13–15^.

SM allows individuals to briefly retain sensory input after a stimulus has ended ^6,7,16–18^, providing the opportunity for this information to be transferred to short-term memory. STM, in contrast, is a limited-capacity storage system that maintains information consciously for short periods of time ^8,9^. A common example of STM in action is remembering a phone number just long enough to write it down.

WM is a more complex and dynamic memory subsystem, originally proposed by Baddeley and Hitch in 1974 and refined in subsequent works ^10,11^. It is defined as a limited-capacity system responsible for the temporary storage and manipulation of information, a process which is necessary for complex tasks such as comprehension, learning, and reasoning. WM has been strongly linked to general cognitive abilities ^19^, with its distinguishing feature being the active manipulation of information, unlike STM which functions primarily as a passive temporary storage system ^8,9^. The development of the WM concept originally arose from the limitations of the modal model of memory, which was prevalent at the time but failed to account for the dynamic nature of cognitive processing. In contrast to this earlier model, the embedded-processes approach ^12^ views WM as an executive system that temporarily activates long-term memories, thereby enabling real-time manipulation of information.

LTM is the subsystem responsible for storing information over extended periods, ranging from hours to months, and in some cases, throughout an individual’s entire life ^20^. LTM encompasses episodic ^21^, procedural ^22^, and semantic memory ^21^, which relate to personal life events, motor skills, and factual knowledge, respectively. Based on the conscious availability of stored information, long-term memories are classified into two main types: procedural (implicit) and declarative (explicit) memory ^23^. Procedural memory underlies a variety of skills, such as learning how to ride a bike, which are performed without conscious effort ^24^. In contrast, declarative memory refers to consciously accessible information that can be actively recalled and verbalized ^24^. Before information can be stored in LTM, it must first be encoded. Encoding is the process of transforming sensory input into a format suitable for long-term storage ^25,26^. This transformation can occur through various modalities, such as visual, acoustic, or semantic, depending on the nature of the information and its associated meanings. Once encoded, information becomes accessible for retrieval or recognition. The process of encoding is closely linked to SM, STM and WM, as information is initially perceived through the senses, briefly held in STM or manipulated by WM, and ultimately encoded into LTM for long-term storage ^27^.

Once a memory is consolidated in LTM, it can be accessed and utilized through two primary methods: recall and recognition ^13^. Recall involves retrieving information from memory, either without any prompts or with cues that are associated with the information ^28^. In contrast, recognition involves identifying specific information as having been previously encountered ^13^. The foundational dual-process framework by Anderson and Bower (1972) ^29^ suggests that recall and recognition are supported by distinct cognitive processes, each with its own set of steps to reach a conclusion. This distinction aligns with findings such as those of Tversky (1973) ^25^, who showed that encoding strategies differ when individuals anticipate a recall test versus a recognition test. When comparing the two, research by Postman et al. (1975) ^13^ demonstrated that performance on recall tests declines more rapidly than on recognition tests. Recent studies have further corroborated this pattern ^30,31^. Within recognition, the literature further differentiates between two distinct processes: familiarity and recollection ^32^. The key difference lies in context dependence—recollection requires a semantic context to properly place the retrieved information, whereas familiarity does not. As a result, recollection tends to exhibit slower forgetting rates compared to familiarity ^33^.

There is also evidence that the mechanisms for recognizing previously encountered information differ from those involved in recognizing novel or unknown stimuli ^34,35^. This distinction is primarily supported by neural evidence, suggesting that the recognition of novelty, or the detection of something unfamiliar, is mediated by processes such as prediction error ^34,36,37^. Recently, we explored this phenomenon through a series of behavioral and neuroscientific studies focused on LTM recognition of a specific type of stimulus: musical sequences. These stimuli are particularly unique because they generate meaning through their combination and progression over time, offering a distinct contrast to the more static stimuli traditionally used in memory research ^36,38,39^. Our findings revealed that individuals could memorize a musical piece with remarkable ease, even after just a few repetitions. Moreover, participants were able to recognize short melodies extracted from the original piece, as well as identify novel melodies or variations of the original sequences. Collectively, these studies suggest that musical sequences offer an effective model for studying memory processes related to sequences that unfold over time and that recognition of novel versus previously memorized sequences is overall more challenging ^34,36^.

Among the various factors that can influence the memory subsystems described above, aging is one of the most well-documented, with numerous studies highlighting that memory is among the cognitive functions most susceptible to decline with advancing age ^40–45^. This is especially true for declarative memory, which tends to deteriorate more rapidly ^46,47^. While this decline is a natural process, certain conditions can accelerate it ^24,43^. Importantly, the effects of aging are not limited to LTM ^45,48^; they also impact WM ^49,50^, which plays a critical role in complex cognitive tasks. Gordon-Salant and Cole (2016) provided evidence ^51^ of this by comparing the speech recognition abilities of younger and older adults, further categorizing them by their WM capacity. This study found that both age and WM capacity independently affect speech recognition, with older adults showing greater difficulty in recognizing sentences, particularly those with lower WM capacity. The interaction between age and WM was significant, suggesting that age-related decline in WM exacerbates difficulties in tasks requiring memory and real-time processing, such as sentence recognition. Interestingly, recent findings indicate that, while aging does not necessarily impair the LTM recognition of familiar musical melodies, older adults perform worse than younger adults when it comes to recognizing variations of these melodies ^36^. This suggests that, although basic recognition abilities may be preserved with age, the ability to adapt to changes or variations in learned sequences declines, potentially due to reduced WM capacity or changes in cognitive flexibility.

While extensive research has been conducted on both WM ^10–12^ and LTM ^13–15^, much remains to be learned about the interaction between the two and the underlying mechanisms that connect them. The two pioneering models of WM, as previously discussed, provide different perspectives on the relationship between WM and LTM. In Baddeley’s model, the central executive component of WM accesses information stored in LTM and uses it to guide present actions ^11^. In contrast, Cowan’s approach conceptualizes WM as a temporary activation of LTM, serving as a “window to the present” ^12^. Despite these differences, both models agree that WM enables individuals to respond to environmental stimuli by integrating incoming information with stored knowledge from LTM. This capacity to manipulate and integrate information is a key reason why WM has been strongly linked to general cognitive abilities ^19^ and why it remains a crucial variable in memory research. Although there is theoretical and empirical support for the interaction between these two memory systems, our understanding of the precise nature of this relationship remains limited.

This is particularly relevant when examining how WM influences various types of recognition and predictive processes ^52,53^. For example, one area of interest is the relationship between WM and the ability to recognize previously memorized information, which depends on the accuracy of predictions about expected outcomes (e.g., the predicted incoming sound in a sequence) compared to the actual outcomes (e.g., the actual incoming sound in the sequence). Similarly, it is relevant to understand how WM relates to the recognition of variations within a sequence, involving prediction error processes (e.g., when the incoming sound in the sequence deviates from the predicted sound). Understanding these dynamics is crucial, especially when dealing with sequential information that evolves over time, where meaning is constructed progressively (as in the example of the sequences of sounds). The gap in our knowledge about how WM interacts with these processes underscores the need for further investigation into how memory systems handle both expected and unexpected changes in sequential patterns.

Building on our extensive research into long-term encoding and recognition of musical sequences ^34,36,54–58^, the current study investigates how WM and LTM interact. We employed musical memory tasks across three distinct datasets, involving 244 healthy participants. This study also examines the effects of aging and musical training on both LTM and WM, drawing on established findings in the field ^59–62^.

## Methods

### Participant samples

We collected demographic and behavioral data from three different samples for a total of 244 participants on three time points: 2021, 2022, and 2024.

The 2021 dataset consisted of 83 participants (49 females) aged 18 to 63 years old (mean age: 28.74 ± 8.10 years). Participants came from Western countries and had homogeneous educational backgrounds. The project was approved by the Institutional Review Board (IRB) of the Center for Music in the Brain, Aarhus University (case number: DNC-IRB-2020-006). Independent, neuroscience results obtained from this dataset have been published in a previous work^36^.

The 2022 dataset comprised 77 participants (43 females) aged 18 to 81 years old (mean age: 45.58 ± 23.31 years). All participants were Danish and had homogenous educational backgrounds. The project was approved by the IRB of the Center for Music in the Brain, Aarhus University (case number: DNC-IRB-2021-012). Previous neuroscientific and behavioral results obtained from this dataset have been published in two previous papers ^36,63^.

The 2024 dataset included 84 participants (52 females) aged 18 to 83 years old (mean age: 45.27 ± 24.17 years). All participants were Danish and had homogenous educational backgrounds. The project was approved by the Ethics Committee of the Central Denmark Region (De Videnskabsetiske Komitéer for Region Midtjylland, ref. 1-10-72-127-23).

The following inclusion criteria were applied to all datasets: (1) normal health (no reported neurological nor psychiatric disease), (2) normal hearing (non-pathological age-related hearing decline was not an exclusion criterium if participants could comfortably perform the task), (3) normal sight or corrected normal sight, (4) individuals aged over 18 years old, and (4) understanding and acceptance of experimental procedure. All experimental procedures complied with the Declaration of Helsinki—Ethical Principles for Medical Research, and informed consent was obtained from all participants before starting the study.

### Long-term memory measure

We employed the old/new auditory recognition task we have used in our previous studies (e.g., ^36,56,63^) to measure long-term memory. The task consists of an encoding and a recognition phase. During the encoding phase, participants listened twice and were asked to memorize as much as possible a shortened version of the right-hand part of J. S. Bach’s Prelude No. 2 in C minor, BWV 847. During the recognition part, participants listened to fragments of the prelude (i.e., memorized sequences) and variations of these fragments (i.e., novel sequences). Memorized sequences consisted of the first five notes from each bar of the prelude, while variations were created by altering the memorized sequences from either the second, third, fourth, or fifth notes. Participants used a response pad and were instructed to press ‘1’ if the displayed sequence was memorized, and ‘2’ if it was novel. We scored responses separately for memorized and for novel sequences. For further details on this task, please see Bonetti et al. (2024) ^36^.

### Working memory measure

Working memory skills were estimated using the Working Memory Index from the Wechsler Adult Intelligence Scale IV ^64^. This measure comprises two subsets: Digit Span, where participants listen to sequences of numbers and reproduce them orally in the same, reversed, or ascending order, and Arithmetic, where they listen to mathematical problems and provide solutions orally without external aids. We combined the raw scores from the Digit Span and Arithmetic subtests to provide a total working memory score.

### Musical training measure

We used the musical training facet from the Goldsmiths Musical Sophistication Index questionnaire ^65^ to measure participants’ history of formal musical training. This part of the questionnaire comprises seven questions related to self-assessed musicianship and extent of musical training and practice. Each item was rated on a seven-point Likert scale, with total scores ranging from 7 (i.e., no history of formal musical training) to 49 (i.e., professional musicians). In the case of datasets from 2022 and 2024, we employed the Danish version of the questionnaire ^66^.

### Statistical analysis

Generalized linear models were used to assess the association between working memory (WM) and long-term memory (LTM) skills. For each of the three datasets, two generalized linear models were computed, one for LTM for memorized sequences and one for LTM for novel sequences, with WM, musical training, and age as predictors. Since measures of memorized and novel LTM were continuous and presented a right-skewed distribution, we used a gamma generalized linear model with a log link function. Akaike Information Criterion (AIC) was used to assess the goodness-of-fit of the models, which was used to compare different model specifications including alternative link functions. Models with the lowest AIC values were used. Statistical analyses were performed using RStudio ^67^.

## Results

Two generalized linear models were computed for each dataset to assess the relationship between the predictors (WM, age, and musical training) and LTM for memorized sequences, and between the predictors and LTM for novel sequences. The relationship between WM and LTM abilities is illustrated in **Figure 1**.

**Figure 1.**
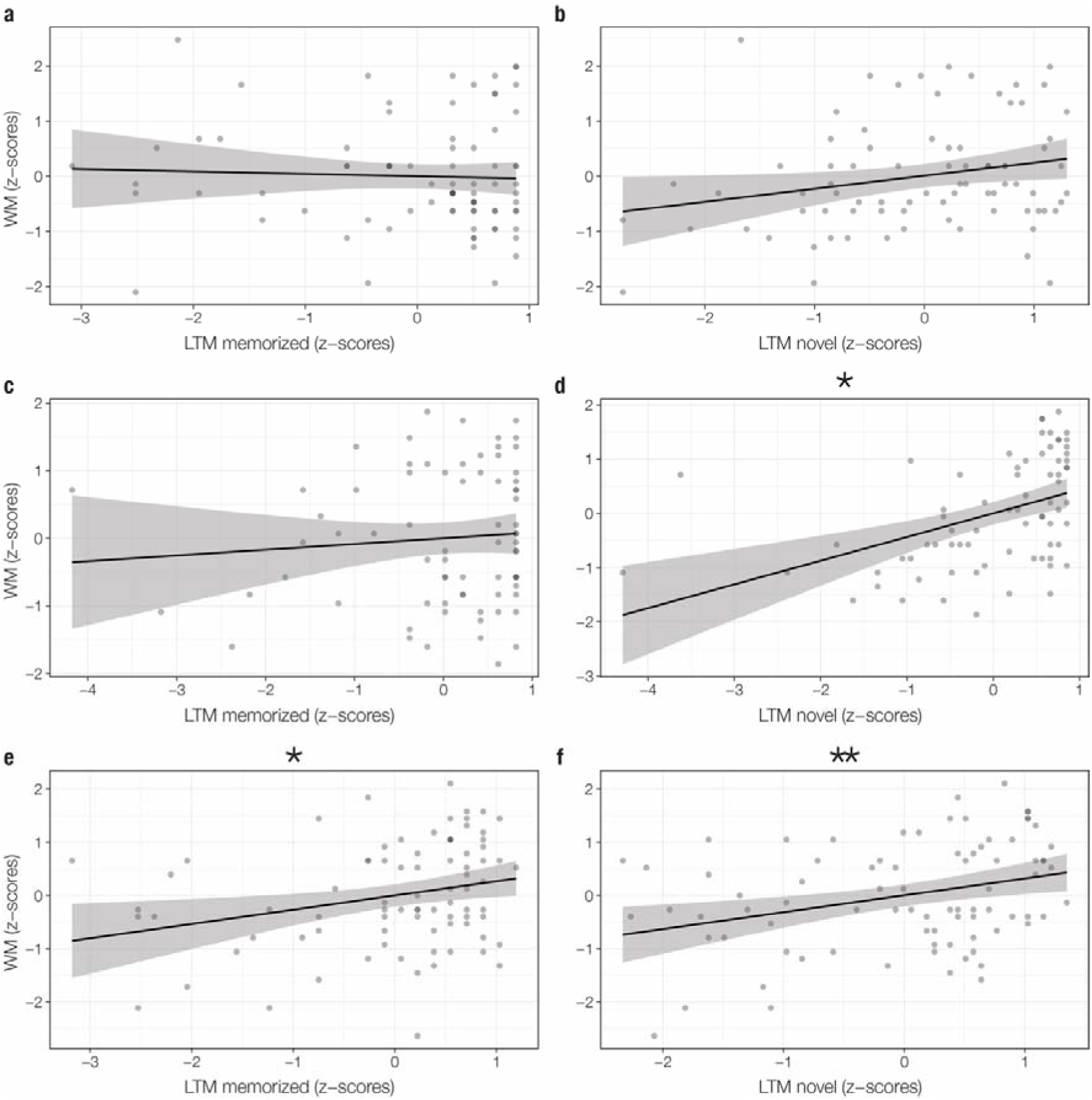
Relationship between working memory (WM) and long-term memory (LTM) abilities. Higher WM scores predict higher LTM scores for categorization of novel sequences in the 2022 (**d**) and 2024 datasets (**f**) and approached the significance for the 2021 dataset (**b**). There was no interaction between WM and LTM for recognition of memorized sequences in the 2021 (**a**) and 2022 (**c**) datasets. However, this was significant for the 2024 dataset (**e**). Note: *p < .05, **p < .01.

In relation to the 2021 dataset, only musical training was significant, contributing to higher scores in LTM for memorized sequences (*B* = .009, *t*-value = 2.703, *p*-value = .009) and LTM for novel sequences (*B* = .008, *t*-value = 2.457, *p*-value = .016), as reported in **Table 1**.

**Table 1.**
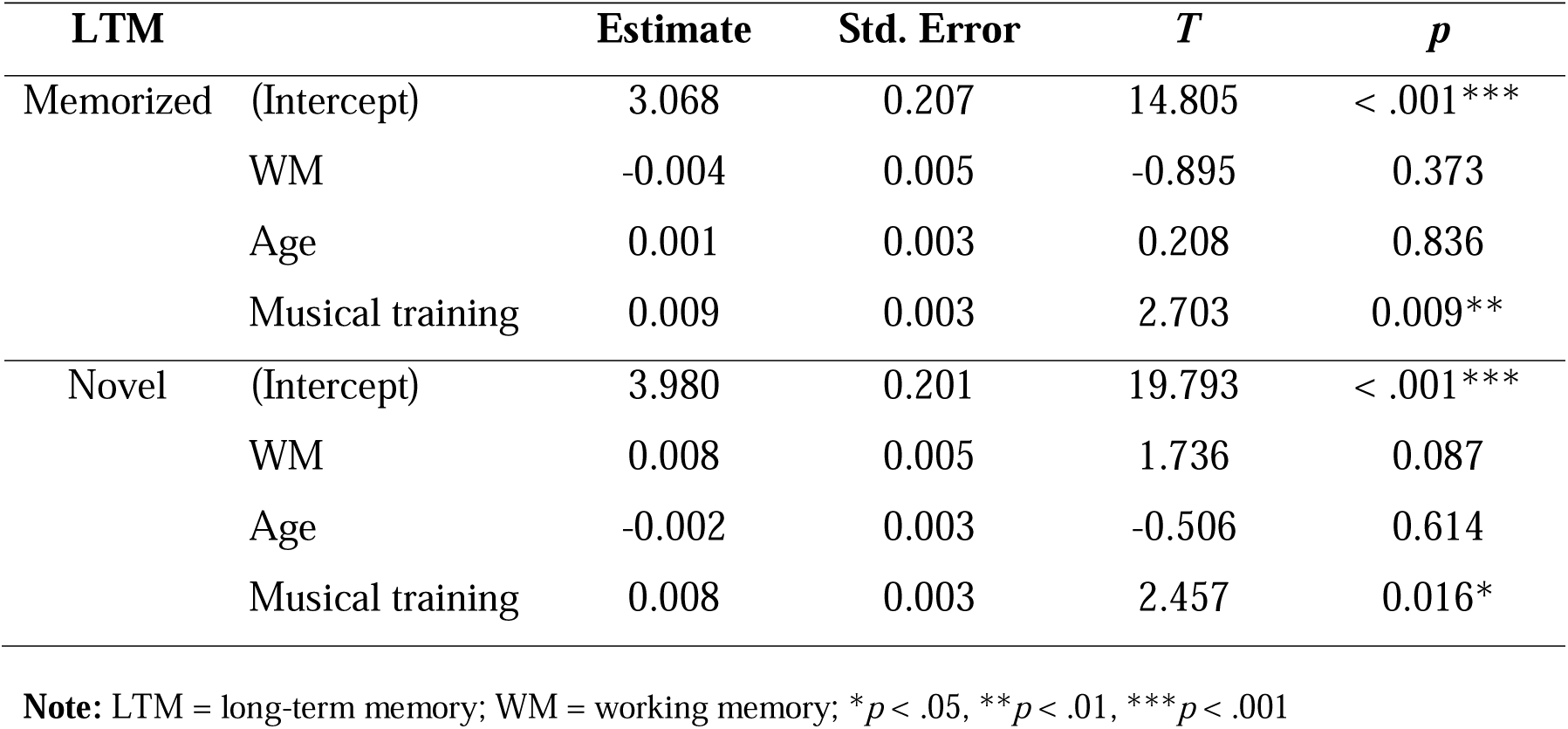
Generalized linear regression models from the 2021 dataset.

Regarding the 2022 dataset, no predictor variables were significantly associated with LTM recognition of memorized sequences. Conversely, the model which included LTM for novel sequences showed that WM contributed to an increase (*B* = .009, *t*-value = 2.620, *p*-value = .011) and age contributed to a decrease (*B* = −.004, *t*-value = −3.133, *p*-value = .003) in LTM for novel sequences. The results for both generalized linear regression models are shown in **Table 2**.

**Table 2.**
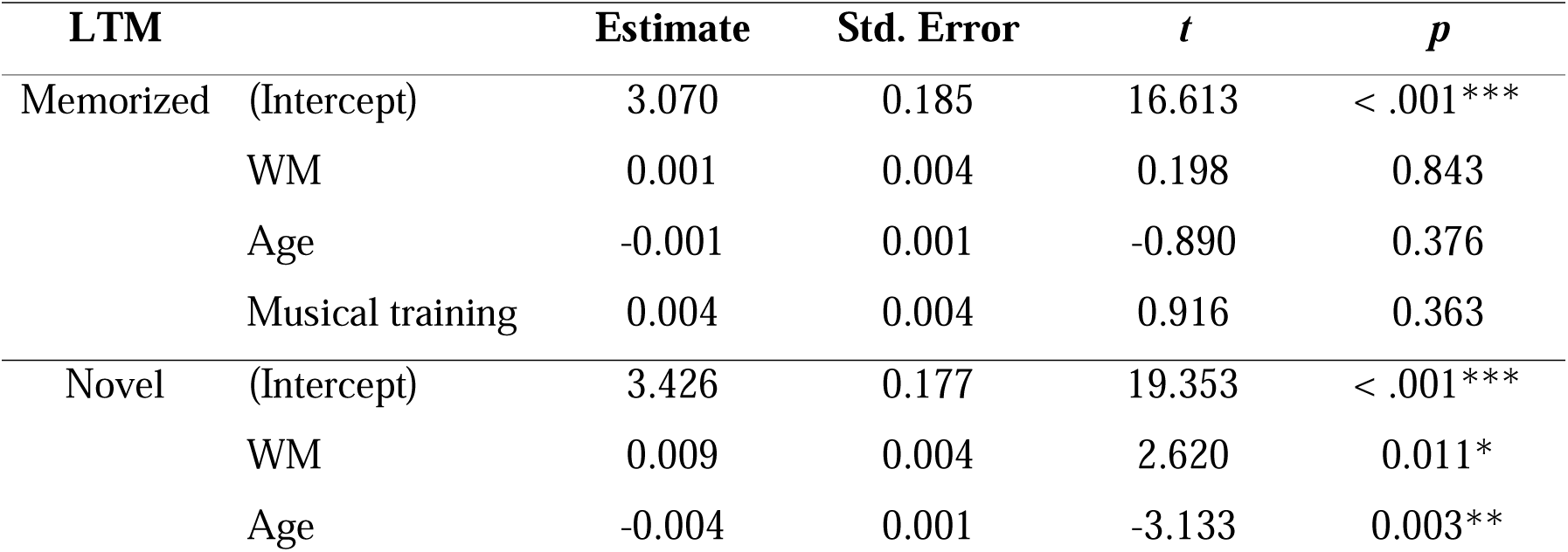
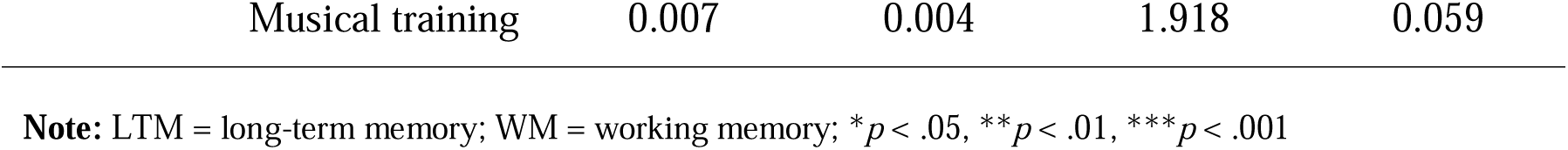
Generalized linear regression models from the 2022 dataset.

**Table 3.** Generalized linear regression models from the 2024 dataset.

| LTM | | Estimate | Std. Error | $t$ | $p$ |
| --- | --- | --- | --- | --- | --- |
| Memorized | (Intercept) | 2.464 | 0.136 | 18.170 | < .001*** |
|  | WM | 0.007 | 0.003 | 2.446 | 0.017* |
|  | Age | 0.000 | 0.001 | -0.449 | 0.655 |
|  | Musical training | 0.005 | 0.003 | 1.617 | 0.110 |
| Novel | (Intercept) | 3.046 | 0.182 | 16.707 | < .001*** |
|  | WM | 0.010 | 0.004 | 2.729 | 0.008** |
|  | Age | -0.003 | 0.001 | -2.424 | 0.018* |
|  | Musical training | 0.004 | 0.004 | 1.003 | 0.319 |

**Note:** LTM = long-term memory; WM = working memory; \* $p < .05$ , \*\* $p < .01$ , \*\*\* $p < .001$

With regards to the 2024 dataset, WM was significant and contributed to higher scores in both LTM for memorized sequences (*B* = .007, *t*-value = 2.446, *p*-value = .017) and for novel sequences (*B* = .010, *t*-value = 2.729, *p*-value = .008). Moreover, older age was associated with lower scores in LTM for novel sequences (*B* = −.003, *t*-value = −2.424, *p*-value = .018).

## Discussion

The results from the 2022 and 2024 datasets revealed a significant relationship between WM and LTM, while the 2021 dataset’s results approached significance. These findings strengthen the concept that memory subsystems are interconnected, aligning with previous models that link WM to LTM ^11,12,15,68^. Notably, this relationship is observed specifically in the identification of novel sequences, suggesting that the role of WM varies between recognizing previously memorized sequences—where predictions are confirmed—and identifying new melodies—where predictions are disrupted, generating prediction errors. This distinction implies that these two LTM processes may operate differently. These findings are consistent with our prior neuroscientific studies, which have shown that distinct neural processes underpin the recognition of memorized versus novel musical sequences ^34,36,54–57,63^. The strong association between WM and the recognition of novel melodies indicates that WM is closely related to prediction error and plays a role in detecting discrepancies. Conversely, WM does not appear to be as involved in the recognition of previously memorized sequences, suggesting that this process might be less complex and may not engage WM to the same extent.

Predictive processes are a fundamental concept in psychology and neuroscience, leading to the formulation of the Predictive Coding Theory (PCT) and extensively reviewed by several authors, including Karl Friston ^52,53,69,70^ and others ^71,72^. In particular, predictions can be confirmed or violated, generating prediction errors in the latter case. Den Ouden et al. (2012) ^73^ offer a comprehensive overview of distinct types of prediction error, outlining their functions and neural underpinnings. They categorize prediction errors into sensory (e.g., oddball paradigm), cognitive (when an event deviates from expectations), and motivational (when value is assigned to an outcome). Despite their different functions and contexts, these types all share a common mechanism of comparing top-down knowledge with bottom-up input. This highlights that prediction error is a core mechanism present across both lower and higher-order brain regions and cognitive processes. Our study aligns with this framework, focusing on cognitive prediction error during the recognition of varied musical sequences. In this task, participants consciously evaluated and predicted the upcoming sounds based on previously heard sequences. The introduction of novel sounds created a cognitive prediction error, as the anticipated sound did not match the actual event. Our findings demonstrate that WM abilities influence performance in tasks involving such cognitive prediction errors, thereby affecting the accuracy of these predictions.

Additional research linking prediction error to WM and LTM underscores the importance of their interaction in memory consolidation and reconsolidation ^74^. In this context, prediction error serves not only as a mechanism for detecting discrepancies but also as a facilitator of learning. Specifically, when there is a mismatch between encoded and experienced information, the process involves activating the memory, destabilizing it through the prediction error, and then updating and reinforcing it. WM plays a crucial role in this process by enhancing the flexibility and efficiency of information updating. This strategy for cognitive updating is supported by several theoretical frameworks, including the PCT ^75,76^ and the Bayesian Brain Hypothesis ^77^, which outline its neural mechanisms and behavioral characteristics.

Our recent studies on the neurophysiology of music encoding and recognition also provide compelling evidence for the relationship between WM and LTM. For example, we demonstrated that WM abilities influence the brain mechanisms involved in encoding single sounds. Specifically, individuals with higher WM exhibited greater recruitment of frontal regions, such as the right frontal operculum, during sound encoding ^54^. In terms of LTM recognition of music, WM was linked to enhanced brain responses when recognizing previously memorized sounds ^57^. This finding was further supported and expanded by our recent work, which showed that WM is associated with the recognition of both previously memorized and varied melodies, with a particularly strong effect for varied melodies. This effect is notably related to the prediction error signal generated by these variations^36^. Interestingly, this result was corroborated by another study in which we found that WM abilities had a more pronounced impact on the prediction error signal compared to age, revealing similar prediction error signals in older adults with high WM and young adults with low WM ^36^. Moreover, our earlier research identified a relationship between enhanced mismatch negativity (MMN)—a brain signal indicative of automatic or sensory prediction error—and higher WM abilities ^78^. This finding is significant as it extends the association between prediction error and WM beyond LTM, implicating a brain signal often interpreted as reflecting automatic sensory memory processes.

In the current study, the relationship between WM and LTM was significantly influenced by participants’ age. Consistent with our recent findings ^36^, we observed that older adults had no significant difficulty recognizing previously memorized melodies, but their ability to recognize varied melodies was notably impaired. In the 2021 dataset, which included mostly younger participants with only a few individuals over 60 years old, the relationship between WM and LTM approached significance but did not reach it. In contrast, in the 2022 and 2024 datasets, which included a substantial proportion of older adults, the relationship between WM and LTM became significant, particularly for LTM related to novel sequences, and exhibited a medium effect size. These results suggest that the association between WM and LTM may become more critical as cognitive decline progresses with age, making tasks like recognizing novel sequences more challenging. For younger participants, the relationship might be masked by a ceiling effect, where WM and LTM are already functioning optimally. In this context, older adults might rely more heavily on their WM resources to compensate for difficulties in recognizing novel sequences, potentially indicating that they allocate more WM resources to handle prediction errors in LTM tasks.

Consistent with our findings, existing literature acknowledges a natural, non-pathological cognitive decline associated with aging, though this decline is not necessarily exacerbated by medical conditions ^42,51^. Studies often highlight episodic (or autobiographical) memory as the primary domain affected by aging, while semantic memory tends to remain relatively intact ^46,47^. Shing et al. ^46^ theorize that this asymmetry leads to difficulties in processing novel information for older adults. This reliance on well-established semantic knowledge might hinder the ability to form new episodic memories efficiently. In our study, the LTM task used, which involves translating sounds into mental (musical) representations, is arguably more aligned with semantic memory than episodic memory. This may explain why older participants did not significantly underperform compared to younger participants in recognizing previously memorized sequences. Additionally, older adults are more susceptible to retroactive interference during recognition tasks, especially when faced with distractors ^41,79^. Their neural activation patterns diverge from those of younger individuals, and they show reduced neural representation of prediction errors in feedback-driven reinforcement learning, although evidence regarding confirmatory prediction abilities remains inconclusive ^80,81^. Our results further support a dissociation between the processes involved in prediction error for novel sequences and the recognition of previously learned information, with this dissociation becoming more pronounced in aging populations. As already mentioned above, this aligns with our recent findings on age-related changes in brain dynamics during music recognition ^36^. We observed that while prediction error signals decrease with healthy aging, recognition of previously memorized sequences is associated with compensatory brain mechanisms, such as increased activity in the left auditory cortex. This increase likely compensates for reduced functioning in memory-related regions of the medial temporal lobe, such as the hippocampus and inferior temporal cortex. The absence of such compensatory mechanisms in older adults corresponds with the reduced behavioral performance observed across datasets.

Finally, our study found a positive association between musical training and memory performance in the 2021 dataset, and this association approached the significance level in the 2022 dataset. This finding supports previous research that associates musical expertise with enhanced auditory, musical, and general cognitive abilities, as well as increased neural responses during these tasks ^60–62,82,83^. For example, Schellenberg demonstrated that even a short formal musical training is linked to higher IQ. In another study, Schulze and Tillmann (2013) ^82^ explored forward and backward recognition of different types of auditory information and found a positive correlation between working memory (WM) for pitch sequences—encompassing both recognition and manipulation of these sequences—and years of musical training. Their findings indicated that increased years of musical training were associated with better performance on tasks involving pitch sequence manipulation, suggesting a higher domain-specific capacity linked to musical expertise. Our own research also supports this association, as demonstrated in a study comparing memory functions in musicians and amateurs versus non-musicians ^61^. However, the underlying reasons for this association remain debated. Schellenberg has recently proposed that the correlation might stem from a higher likelihood of individuals with higher IQs engaging in musical training, rather than musical training directly enhancing cognitive abilities ^60^. This challenges the earlier hypothesis that musical training itself improves cognitive functions. While further research is needed to clarify this relationship, our current study contributes confirmatory evidence of the association between musical expertise and memory abilities.

To conclude, our findings reveal that WM capacity is significantly associated with the identification of novel sequences but is not similarly related to the recognition of previously memorized sequences. This suggests that different cognitive processes are involved in handling confirmatory predictions versus prediction errors. Benefitting from large sample sizes and replication across multiple datasets, our study significantly advances the understanding of the relationship between WM and LTM, highlighting how these subsystems are influenced by aging and musical training. However, as our study primarily identifies positive relationships between different memory subsystems, further research is needed to elucidate the specific mechanisms through which WM arguably impacts LTM processes. Furthermore, exploring this association in populations with memory impairments, such as individuals with dementia or significantly reduced cognitive abilities, could yield additional valuable insights.

## Data availability

The data will be made available upon reasonable request.

## Code availability

The code used for computing descriptive statistics and plotting results is freely available on GitHub: https://github.com/gemmaferu/interactions-ltm-wm.

## Acknowledgements

The Center for Music in the Brain (MIB) is funded by the Danish National Research Foundation (project number DNRF117).

LB is supported by Lundbeck Foundation (Talent Prize 2022), Carlsberg Foundation (CF20-0239), Center for Music in the Brain, Linacre College of the University of Oxford, Society for Education and Music Psychology (SEMPRE’s 50th Anniversary Awards Scheme), and Nordic Mensa Fund.

GFR is supported by the Center for Music in the Brain and Nordic Mensa Fund.

MLK is supported by the Center for Music in the Brain and Centre for Eudaimonia and Human Flourishing, which is funded by the Pettit and Carlsberg Foundations.

ES is supported by the Luxembourg National Research Fund (FNR) (17906488)

Additionally, we thank Fundación Mutua Madrileña (Mutua Madrileña Foundation, Madrid, Spain) for the support provided to GFR and Mensa: The International High IQ Society (Italian section) for the support provided to FC.

## Author contributions

LB and GFR conceived the hypotheses. LB, GFR, EB, and MLK designed the study. LB, GFR, MLK, EB, and PV recruited the resources for the experiment. LB, GFR, ERO, FC, ES, MK, MAH, and LF collected the data. GFR performed statistical analysis, with the contribution of LB. EB, MLK and PV provided essential help to interpret and frame the results within the psychological and neuroscientific literature. LB, GFR, and SSA wrote the first draft of the manuscript. GFR prepared the figures. All the authors contributed to and approved the final version of the manuscript.

